# Genotype, antimicrobial resistance and virulence profiles of livestock-derived enterotoxigenic *Escherichia coli* (ETEC) in the United States, 1970-2023

**DOI:** 10.1101/2023.12.04.570031

**Authors:** Yezhi Fu, Erin M. Nawrocki, Nkuchia M. M’ikanatha, Edward G. Dudley

**Affiliations:** School of Agriculture, Shenzhen Campus of Sun Yat-sen University, Shenzhen, China; Department of Food Science, The Pennsylvania State University, University Park, PA, USA; Pennsylvania Department of Health, Harrisburg, PA, USA; E. coli Reference Center, The Pennsylvania State University, University Park, PA, USA

**Author notes:** Corresponding authors: (Y.F) or (E.G.D).

## Abstract

Enterotoxigenic *Escherichia coli* (ETEC) is a significant pathogen in both cattle and pigs, causing diarrhea in these animals and leading to economic losses in the livestock industry. Understanding the dissimilarity in genotype, antimicrobial resistance (AMR), and virulence between bovine and swine ETEC is crucial for development of targeted preventive and therapeutic approaches for livestock. However, a comprehensive study on this area remains lacking. Here, we performed whole-genome sequencing-based analyses of bovine (*n* = 554) and swine (*n* = 623) ETEC collected in the US over a 53-year period. We identified distinct ETEC genotypes (*fimH* type, O antigen, H antigen, sequence type) in cattle and pigs. Further, specific AMR and virulence profiles were associated with bovine and swine ETEC. Compared to swine ETEC, bovine ETEC were less diverse in genotypes, had a significantly (*p* < 0.001) lower number of AMR genes per isolate but higher co-occurrence of Shiga toxin and enterotoxin genes. Our results provide an overview of the key genomic differences between bovine and swine ETEC in the US, which might be attributed to host adaptation and antibiotic usage practice. Ongoing surveillance and research are essential to monitor the genetic diversity and AMR patterns of ETEC in different host species.

## Introduction

Enterotoxigenic *Escherichia coli* (ETEC) are a pathovar of *E. coli* species that can produce heat-stable (ST) and/or heat-labile (LT) enterotoxin in the small intestine of humans or livestock^1, 2^. The enterotoxins stimulate the host’s intestine to secrete fluid, thus leading to diarrhea. ETEC are a major enteric pathogen that account for diarrhea among children under five years old in the developing world, responsible for an estimated 84.4 million diarrhea episodes and 44,400 deaths in 2015^3^. The bacterial pathogen is also one of the most common causes of diarrhea outbreaks in young animals, particularly in calves and piglets^4^. The disease caused by ETEC is also known as neonatal diarrhea in calves/piglets and post-weaning diarrhea (PWD) in piglets^4^. Clinical signs of ETEC infection in calves and piglets include watery diarrhea, dehydration, depression, anorexia, and fever. Calves and piglets that are affected with ETEC can become severely dehydrated and lose weight rapidly, which can be life-threatening if not treated promptly^5^. Due to reduced productivity, treatment costs, and animal welfare concerns, ETEC-associated diarrhea represents one of the most economically important diseases in the livestock industry.

Although bovine and swine ETEC strains belong to the same species, *Escherichia coli*, they can exhibit differences in their genetic makeup. A comprehensive understanding of the dissimilarity in genotype, antimicrobial resistance (AMR) and virulence of bovine and swine ETEC can be useful in developing targeted interventions and vaccines to prevent and control ETEC infections in livestock. Previous studies have already differentiated human and swine ETEC based on the specific enterotoxins and colonization factors produced by the pathovar^6–8^. For example, human ETEC are more likely to carry colonization factors such as CFA/I, CFA/II (CS1, CS2, CS3), and CS6^7^, while swine ETEC typically carry colonization factors like F4 (K88), F5 (K99), F6 (987P), and F18^6^. Further, the ST enterotoxin produced by human and swine ETEC usually belongs to subtype STaH and StaP, respectively^8^. However, the two virulence factors alone are unable to differentiate bovine and swine ETEC as both share the common enterotoxins and colonization factors^4^.

While the enterotoxins and colonization factors in bovine and swine ETEC could be the same, other factors (e.g., *fimH* type, O antigen/H antigen, sequence type, AMR, virulence factors such as Shiga toxins) may vary. The dissimilarities between bovine and swine ETEC strains are subject to change over time due to various factors, including host environments, genetic mutations, horizontal gene transfer (HGT), and changes in agricultural practices and antibiotic use. For instance, due to the different antibiotic use in farming practices, specific resistance profiles and prevalence of resistant strains may differ between host species^9^. Further, bacterial pathogens may evolve to thrive in their specific host environments, and this host-specific adaptation can be reflected in their genotype, virulence factors, and ability to cause disease^10, 11^. As new ETEC strains with different AMR and virulence factor profiles may emerge over time among different host species, proper identification and differentiation of these ETEC strains are critical for implementing effective prevention and control measures. However, research in this area is scarce and ongoing research using advanced techniques, such as whole-genome sequencing (WGS) may help us gain new insights into the genetic diversity of ETEC from different host species.

WGS is a powerful technique that can provide detailed information about pathogen genomics^12, 13^, allowing for the analysis of the entire genetic content of ETEC strains. In this study, we performed WGS-based subtyping and analyses on a collection of ETEC isolates collected from bovine (*n* = 554) and swine (*n* = 623) hosts during 1970-2023 in the US. The overall goal of this study is to understand the genetic dissimilarities between bovine and swine ETEC in the US. By comparing the genomes of bovine and swine ETEC using WGS-based analyses, our specific objectives are to: 1) identify the major genotypes (e.g., *fimH* type, O and H serogroups, sequence type) of ETEC circulating in bovine and swine hosts; 2) monitor the AMR patterns and trends of ETEC in bovine and swine ETEC; 3) characterize the specific virulence factors in bovine and swine ETEC. As variations in host ecology such as host diet, antibiotic usage, and host density can have a profound effect on pathogen adaptation^10^, we expect that distinct variants of ETEC with specific genetic characteristics (e.g., genotypes, AMR patterns, virulence factors) may emerge in bovine and swine hosts.

## Results

### Collection of ETEC from bovine and swine hosts

A total of 1,177 ETEC isolates from US bovine and swine hosts were retrieved from EnteroBase on March 10, 2023 (Supplementary Data 1). Of note, among the 1,177 genomes at EnteroBase, our group sequenced and uploaded 477 genomes deposited under BioProject <u>PRJNA357722</u> (Supplementary Data 1), accounting for 40.5% of the whole collection. In the ETEC collection, 554 ETEC isolates had a bovine origin, which were collected in 28 US states between 1976 and 2023 (Fig. 1a), and 623 ETEC isolates had a swine origin, which were collected in 35 US states between 1970 and 2023 (Fig. 1b). The bovine ETEC mainly came from California, Nebraska, and Texas, while the swine ETEC were primarily from Pennsylvania, South Dakota, Minnesota, and Iowa. The geographical distribution of the bovine and swine ETEC isolates (Fig. 1) appeared to reflect the main bovine and swine farming states in the US^14^. It should be noted that the focus of this study is on bovine and swine ETEC strains originating from the US. The rationale behind this emphasis lies in the scarcity of sequencing data pertaining to livestock ETEC isolates from other countries. For example, only ∼70 bovine ETEC isolates outside the US were available at EnteroBase as of the retrieval time. **Bovine and swine ETEC having distinct predominant genotypes.** We performed *in silico* genotyping of the bovine and swine ETEC genomes using ClermonTyper for Clermont phylogroup type^15^, FimTyper for *fimH* type^16^, 7-gene multilocus sequence typing (MLST) for sequence type (ST)^17, 18^, and ECTyper^19^ in combination with EtoKi EBEis (EnteroBase *Escherichia in silico* serotyping module from EnteroBase Tool Kit)^20^ for serotype prediction (O and H serogroups). Generally, distinct genotypes regarding Clermont phylogroup, *fimH*, ST, and serotype predominated in bovine and swine ETEC (Fig. 2). Compared to bovine ETEC, swine ETEC were more diverse in terms of genotypes. Detailed information on genotypes was provided as follows and in Supplementary Data 2:

**Fig. 1:**
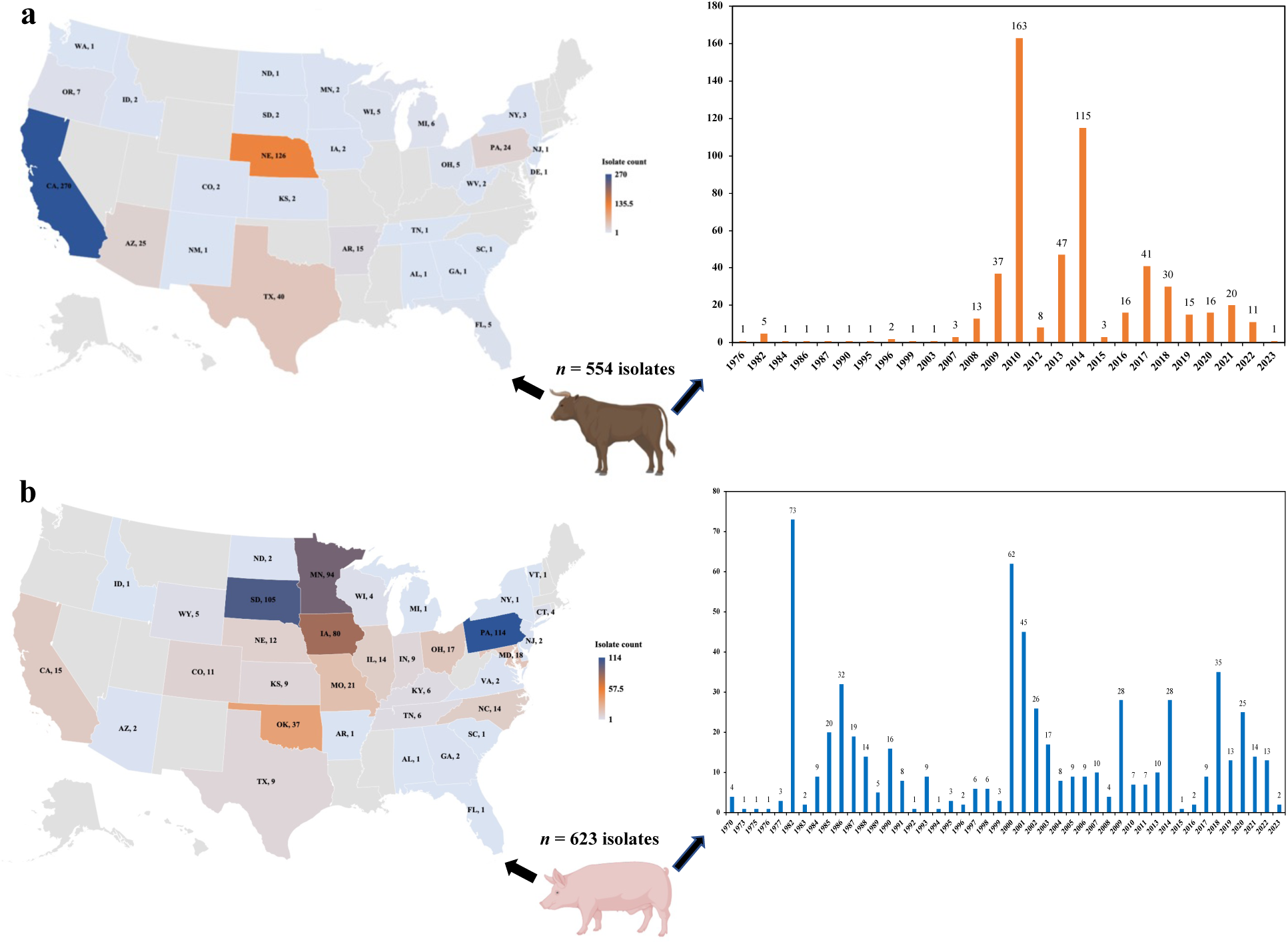
Enterotoxigenic *E. coli* (ETEC) from bovine and swine hosts used in this study. **a**, Geographical distribution and collection years of bovine ETEC isolates originating from the US. **b**, Geographical distribution and collection years of swine ETEC isolates originating from the US.

**Fig. 2:**
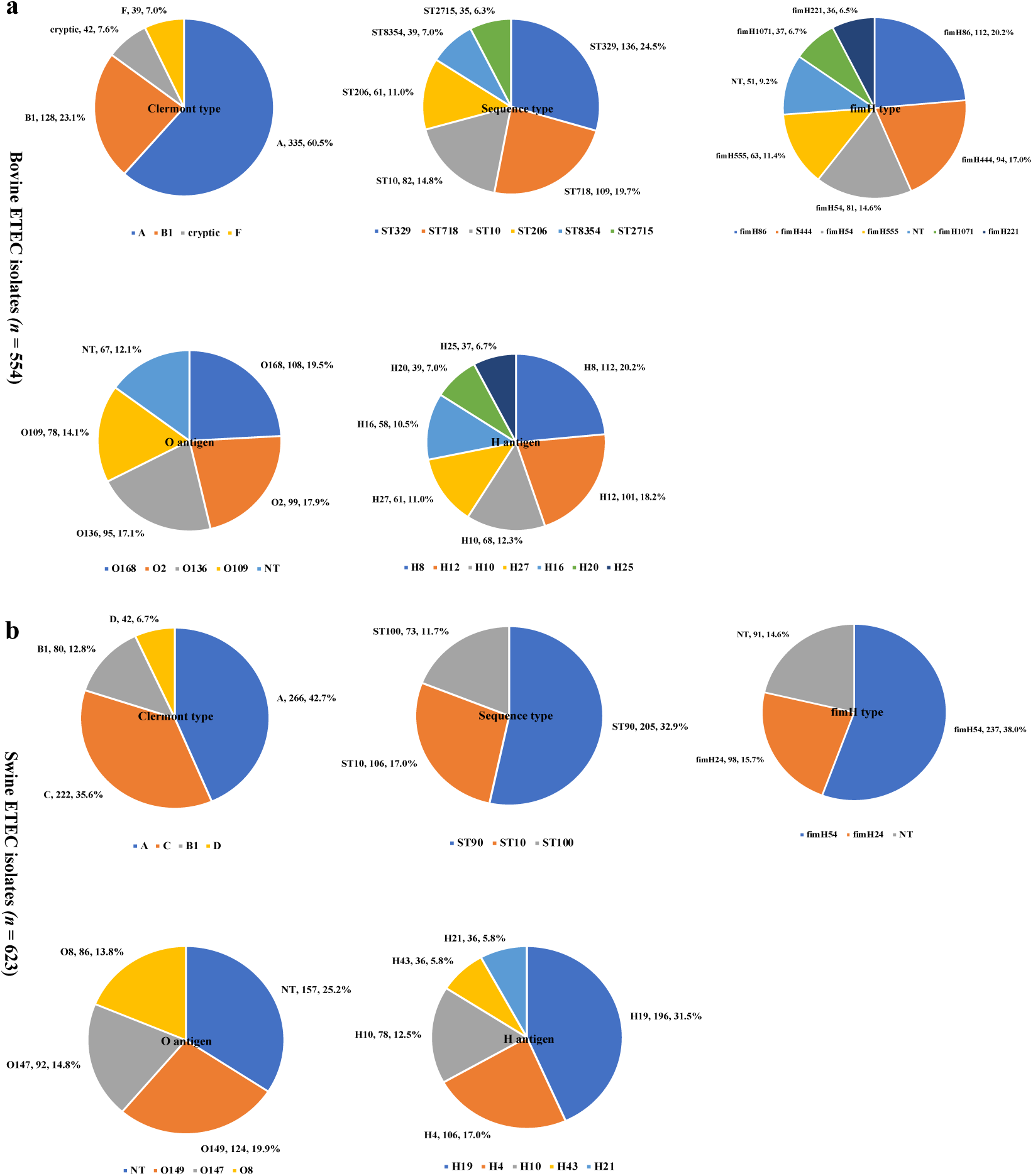
Genotypes (Clermont phylogroup, sequence type, *fimH* type, O and H serogroups) of bovine and swine ETEC in the US. **a**, Genotypes of bovine ETEC. **b**, Genotypes of swine ETEC. The pie charts only show the individual genotype that accounts for more than 5.0% of each collection. Number and percentage following a genotype name in **a** and **b** indicate the number and percentage of the bovine or swine ETEC isolates identified as the specific genotype. “NT” represent “nontypeable”. The full genotype information of the bovine and swine ETEC isolates can be found in Supplementary Data 2.

Bovine ETEC were distributed in six Clermont phylogroups [Clermont type A (60.5%), B1 (23.1%), cryptic (7.6%), F (7.0%), E (1.1%), and C (0.7%)], while a total of seven Clermont phylogroups [Clermont type A (42.7%), C (35.6%), B1 (12.8%), D (6.7%), E (1.0%), F (0.8%), and cryptic (0.3%)] were found in swine ETEC. Clermont phylogroup D was only detected in swine ETEC. In addition, although Clermont phylogroup C accounted for 35.6% (222/623) of the swine ETEC isolates, it only represented 0.7% (4/554) of the bovine ETEC isolates. It is also noteworthy that a considerable number (7.6%) of bovine ETEC isolates belonged to the cryptic phylogroup, while only 0.3% of the swine ETEC were in this phylogroup.

The STs of bovine ETEC were much less diverse than those of swine ETEC. Specifically, a total of 36 STs were identified in the 554 bovine ETEC isolates, with ST329 (24.5%), ST718 (19.7%), ST10 (14.8%), ST206 (11.0%), ST8354 (7.0%), and ST2715 (6.3%) accounting for > 80.0% of the total bovine ETEC collection. On the other hand, a total of 85 STs were detected in the 623 swine ETEC isolates. The dominant STs in swine ETEC were limited to ST90 (32.9%), ST10 (17.0%), and ST100 (11.7%), and all other STs were less than 5.0% of the total swine ETEC collection.

Similar to STs, the diversity of *fimH* type was much lower in bovine ETEC than in swine ETEC. A total of 17 and 36 *fimH* types were identified in bovine and swine ETEC, respectively. The predominant *fimH* types were *fimH*86 (20.2%), *fimH*444 (17.0%), *fimH*54 (14.6%), *fimH*555 (11.4%), *fimH*1071 (6.7%), and *fimH*221 (6.5%) in bovine ETEC (*n* = 554), while the most prevalent *fimH* types were *fimH*54 (38.0%) and *fimH*24 (15.7%) in swine ETEC (*n* = 623). All the other *fimH* types in bovine and swine ETEC accounted for less than 5.0% of each collection. Notably, 9.2% of bovine ETEC and 14.6% of the swine ETEC were not typeable via FimTyper.

A large variation of O and H serogroups was recorded in our serogroup prediction. In specific, a total of 21 O serogroups and 20 H serogroups were detected in the 554 bovine ETEC isolates, with O168 (19.5%), O2 (17.9%), O136 (17.1%), and O109 (14.1%) being the dominant O groups and H8 (20.2%), H12 (18.2%), H10 (12.3%), H27 (11.0%), H16 (10.5%), H20 (7.0%), and H25 (6.7%) being the major H groups. On the other hand, the 623 swine ETEC were represented by 28 O serogroups and 32 H serogroups, among which O149 (19.9%), O147 (14.8%), and O8 (13.8%) are the main O groups and H19 (31.5%), H4 (17.0%), H10 (12.5%), H43 (5.8%), and H21 (5.8%) are the prevailing H groups. It is noteworthy that the nontypeable O antigens accounted for 12.1% (67/554) of the bovine ETEC isolates, and 25.2% (157/623) of the swine ETEC isolates. However, all the bovine ETEC isolates were assigned H serogroups, and only 2.1% (13/623) of the swine ETEC isolates did not have identifiable H serogroups.

### Significantly higher number of AMR genes present in swine ETEC than in bovine ETEC

AMR profiling via AMRFinder^21^ detected 95 types of AMR genes or their variations among the 1,177 ETEC isolates, conferring resistance to 14 antibiotic classes (i.e., aminoglycoside, beta-lactam, bleomycin, chloramphenicol, colistin, fluoroquinolone, fosfomycin, lincosamide, macrolide, rifamycin, streptothricin, sulfonamide, tetracycline, and trimethoprim) (Supplementary Data 3). Among the detected AMR genes, four were solely found in bovine ETEC isolates (i.e., *bla*_CTX-M-1_, *bla*_CTX-M-27_, *bla*_CTX-M-32_, and *dfrA23*), 51 were unique to swine ETEC, and 40 were identified both in bovine and swine ETEC (Fig. 3a). Only eight AMR genes were carried by more than 5.0% of the bovine ETEC isolates (Fig. 3b), i.e., *blac*_EC_ (50.9%), *blac*_EC-18_ (22.6%), *blac*_EC-15_ (16.6%), *tet*(A) (12.1%), *blac*_EC-8_ (9.2%), *sul1* (7.0%), *aph(3′′)-Ib* (6.1%), and *aph(6)-Id* (6.0%), which conferred resistance to beta-lactam, tetracycline, sulfonamide, and aminoglycoside. However, 27 AMR genes were harbored by higher than 5.0% of the swine ETEC isolates (Fig. 3c), conferring resistance to tetracycline, beta-lactam, aminoglycoside, sulfonamide, chloramphenicol, bleomycin, trimethoprim. The top ten detected AMR genes in swine ETEC were *tet*(B) (53.1%), *tet*(D) (48.2%), *blac*_EC_ (41.9%), *blac*_EC-13_ (37.6%), *aph(3′′)-Ib* (37.2%), *aph(6)-Id* (37.2%), *aph(3′)-Ia* (34.3%), *tet(A)* (32.7%), *sul2* (29.4%), and *blac*_TEM-1_ (27.8%).

**Fig. 3:**
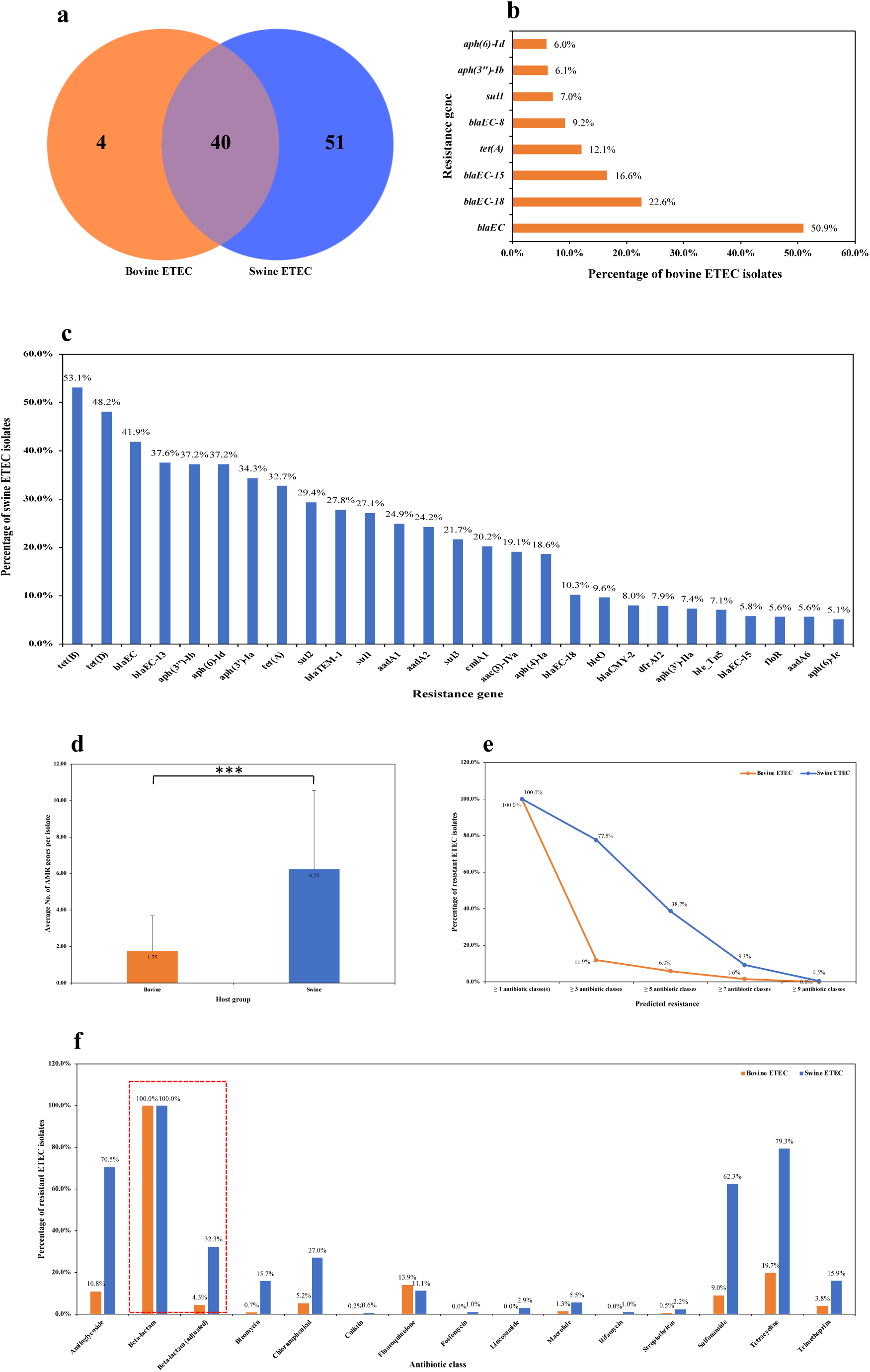
Antimicrobial resistance (AMR) profiles of ETEC from cattle and pigs in the US. **a**, Number of AMR genes unique to or shared by bovine and swine ETEC. **b**, Percentage of bovine ETEC isolates positive for a specific resistance gene. **c**, Percentage of swine ETEC isolates positive for a specific resistance gene. **d**, Average number of AMR genes per isolate detected in bovine and swine ETEC. **e**, Percentage of bovine or swine ETEC isolates predicted to be resistant to ≥ 1, 3, 5, 7, and 9 antibiotic classes. **f**, Percentage of bovine or swine ETEC isolates predicted to be resistant to individual antibiotic classes. Only the individual resistance genes accounting for > 5.0% of each collection are shown in **b** and **c**. The *** in **d** indicates a significance level (*p* value) < 0.001. In **f**, the AMR genes in the antibiotic class “beta-lactam” include all the beta-lactam resistance genes detected in this study, while the AMR genes in the antibiotic class “beta-lactam (adjusted)” include all the beta-lactam resistance genes except the *blac*_EC_ family genes as genes in this family do not confer phenotypic resistance.

The average number of AMR genes per Isolate carried by bovine ETEC was less than 2, which was significantly (*p* < 0.001) lower than that carried by swine ETEC (Fig. 3d; > 6 AMR genes per isolate). Further, the percentage of bovine ETEC isolates predicted to be resistant to ≥ 1, 3, 5, 7, and 9 antibiotic classes was 100.0%, 11.9%, 6.0%, 1.6%, and 0.0%, respectively, while it was 100.0%, 77.5%, 38.7%, 9.3%, and 0.5% for swine ETEC isolates, respectively (Fig. 3e). We also found that the percentage of swine ETEC isolates predicted to be resistant to individual antibiotic classes was mostly much higher than that of bovine ETEC isolates (swine ETEC vs bovine ETEC: beta-lactam 100.0% vs 100.0%, tetracycline 79.3% vs 19.7%, aminoglycoside 70.5% vs 10.8%, sulfonamide 62.3% vs 9.0%, chloramphenicol 27.0% vs 5.2%, trimethoprim 15.9% vs 3.8%, bleomycin 15.7% vs 0.7%, fluoroquinolone 11.1% vs 13.9%, macrolide 5.5% vs 1.3%, streptothricin 2.2% vs 0.5%, colistin 0.6% vs 0.2%) (Fig. 3f). Moreover, predicted resistance to fosfomycin, lincosamide, or rifamycin was only detected in swine ETEC, not in bovine ETEC (Fig. 3f).

### Genetic determinants of AMR to beta-lactam and fluoroquinolone in bovine and swine ETEC

Although the overall percentage of predicted resistant isolates was overwhelmingly higher in swine ETEC than in bovine ETEC, the percentage of bovine and swine ETEC isolates predicted to be resistant to beta-lactam (bovine ETEC: 100.0% vs swine ETEC: 100.0%) or fluoroquinolone (bovine ETEC: 13.9% vs swine ETEC: 11.1%) was at similar levels (Fig. 3f). A close look into the genetic determinants of AMR to beta-lactam identified that 100.0% (554/554) of the bovine ETEC isolates (Fig. 4a) and 99.7% (621/623) of the swine ETEC isolates (Fig. 4b) carried a *blac*_EC_ family gene (i.e., *blac*_EC_, *blac*_EC-8_, *blac*_EC-13_, *blac*_EC-13_, *blac*_EC-15_, *blac*_EC-18_, or *blac*_EC-19_). It should be noted that the *blac*_EC_-associated beta-lactam resistance genes detected in this study were frequently found in genomes of beta-lactam susceptible *E. coli* isolates^22^, suggesting that AMR genes of this family typically had no effect on phenotypic resistance. However, the *blac*_EC_ family genes can be activated to confer phenotypic resistance in the presence of *ampC* promoter mutations^23^. Based on the information, we recalculated the percentage of beta-lactam resistant isolates in bovine ETEC and swine ETEC by excluding the *blac*_EC_ family genes without a point mutation in *ampC* promoter. The adjusted percentage of predicted beta-lactam resistant isolates was 4.3% (24/554) in bovine ETEC, and 32.3% (201/623) in swine ETEC, respectively (Fig. 3f). Further, the predicted beta-lactam resistance of bovine ETEC were attributed to extended-spectrum beta-lactam (ESBL) resistance genes such as *bla*_CTX-M-1_ (0.2%), *bla*_CTX-M-27_ (0.2%), *bla*_CTX-M-32_ (0.2%), *bla*_OXA-1_ (0.4%), and *bla*_OXA-2_ (0.4%), or beta-lactam resistance genes such as *bla*_TEM-1_ (2.9%) and *bla*_CMY-2_ (0.7%) (Fig. 4a). On the other hand, the predicted beta-lactam resistance of swine ETEC was primarily due to the presence of beta-lactam resistance genes such as *bla*_TEM-1_ (27.8%) and *bla*_CMY-2_ (8.0%), and occasionally ESBL genes such as *bla*_CARB-2_ (0.2%), *bla*_OXA-1_ (0.3%), *bla*_OXA-2_ (0.6%), *bla*_SHV-12_ (0.5%), *bla*_TEM-150_ (0.2%), and *bla*_TEM-217_ (0.3%) (Fig. 4b). Notably, 2.4% (15/623) of the swine ETEC harbored a point mutation in *ampC* promoter (T32A), which was not detected in bovine ETEC. In summary, bovine and swine ETEC showed different levels of predicted resistance to beta-lactam; however, ESBL genes were not commonly detected in both ETEC isolates.

**Fig. 4:**
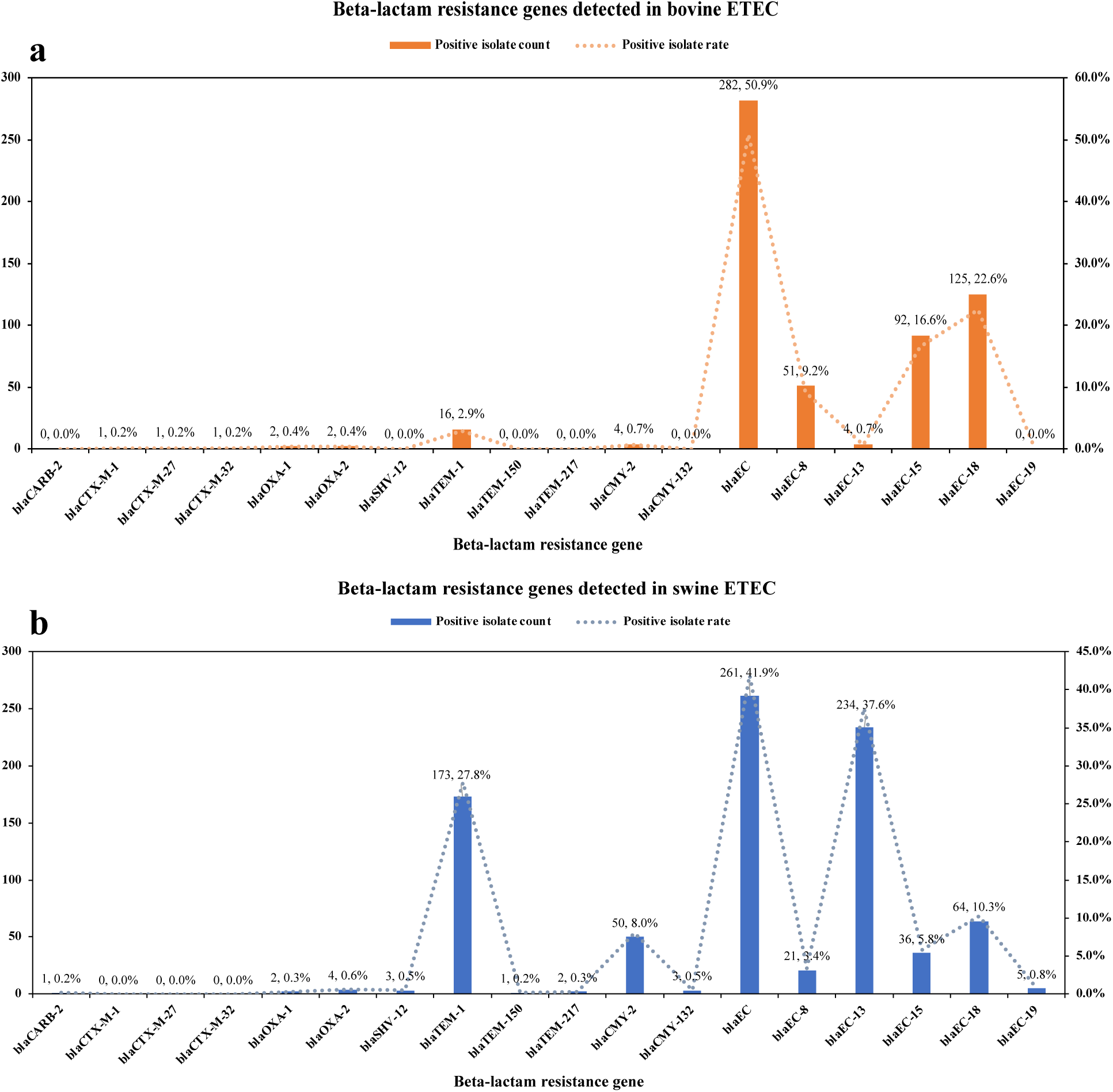
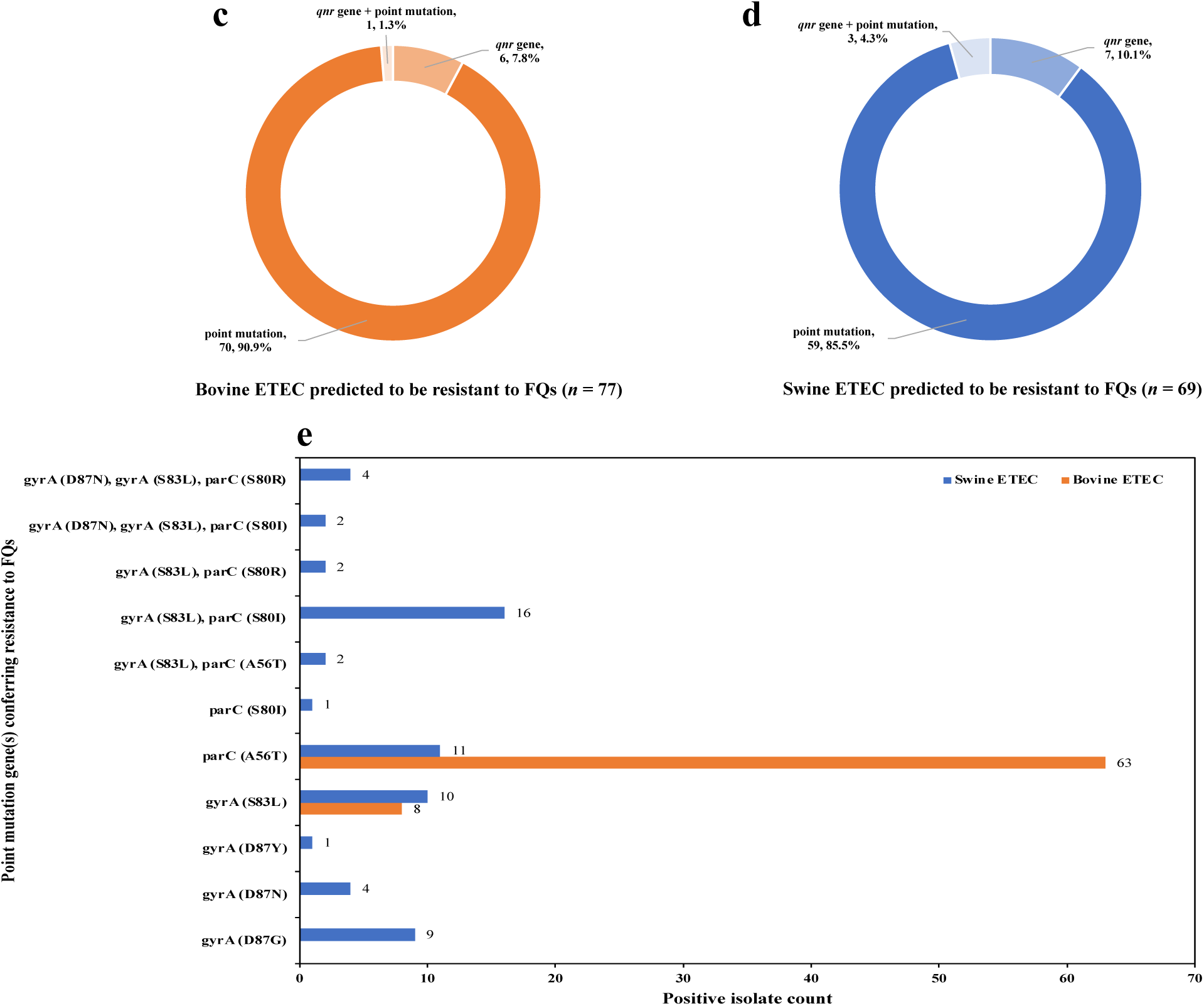
Genetic determinants of antimicrobial resistance to beta-lactams and fluoroquinolones (FQs) in bovine and swine ETEC. **a**, Beta-lactam resistance genes detected in bovine ETEC. **b**, Beta-lactam resistance genes detected in swine ETEC. **c**, Number and percentage of bovine ETEC isolates positive for point mutation, *qnr* gene, or their combination that confer resistance to FQs. **d**, Number and percentage of swine ETEC isolates positive for point mutation, *qnr* gene, or their combination that confer resistance to FQs. **e**, Point mutation genes conferring resistance to FQs detected in bovine and swine ETEC isolates.

A combination of AMRFinder and PointFinder^24^ was used to determine the genetic determinants of AMR to fluoroquinolone (FQ). For the 77 bovine ETEC isolates predicted to be resistant to FQ, 90.9% (70/77) had a point mutation in *gyrA* or *parC* gene, 7.8% (6/77) carried a plasmid-mediated quinolone resistance (PMQR) gene *qnr*, and 1.3% (1/77) presented both a point mutation and a *qnr* gene (Fig. 4c). Similarly, the predicted resistance to FQ in swine ETEC (*n* = 69) was mainly due to point mutations in *gyrA* or *parC* (85.5%, 59/69), followed by the carriage of *qnr* (10.1%, 7/69), and a co-occurrence of point mutation and *qnr* (4.3%, 3/69) (Fig. 4d). Interestingly, although both bovine and swine ETEC isolates exhibited FQ resistance primarily due to point mutations in the quinolone resistance-determining region (QRDR), there were differences in the types and combinations of point mutations. In bovine ETEC isolates, point mutations were either *gyrA* (S83L) or *parC* (A56T), and each isolate contained only one type of point mutation (Fig. 4e). In swine ETEC isolates, however, a more diverse range of point mutation types was observed, including *gyrA* (D87G), *gyrA* (D87N), *gyrA* (D87Y), *gyrA* (S83L), *parC* (A56T), and *parC* (S80I). Co-occurrence of two or three types of point mutation was observed in 26 swine ETEC isolates [e.g., *gyrA* (S83L) + *parC* (A56T), *gyrA* (D87N) + *gyrA* (S83L) + *parC* (S80I)] (Fig. 4e). It is well documented that multiple point mutations confer higher resistance to FQ than a single point mutation^25^, indicating that the swine ETEC isolates with a combination of target-site gene mutations may be more resistant to FQ than the swine and bovine ETEC isolates that only carried one point mutation gene.

### Positive correlation between plasmid replicon content and AMR gene prevalence in bovine and swine ETEC

Plasmid profiling via PlasmidFinder^26^ identified a total of 52 types of plasmid replicons in bovine and swine ETEC (Supplementary Data 4). Among these plasmid replicon types, two were unique to bovine ETEC, 20 were specific to swine ETEC, and 30 were detected in both bovine and swine ETEC (Fig. 5a). The top five plasmid replicons detected in bovine ETEC were IncFIB(AP001918)_1 (88.8%; 492/554), Col156_1 (35.6%; 197/554), ColRNAI_1 (33.6%; 186/554), IncI1_1_Alpha (16.1%; 89/554), and Col(MG828)_1 (13.9%; 77/554); for swine ETEC, they were ColRNAI_1 (82.5%; 514/623), IncFIB(AP001918)_1 (77.7%; 484/623), IncFIC(FII)_1 (53.5%; 333/623), IncI1_1_Alpha (50.1%; 312/623), and IncFII(pSE11)_1_pSE11 (40.4%; 252/623). Similar to the average number of AMR genes per isolate, the average number of plasmid replicons per isolate in swine ETEC (six plasmid replicons per isolate) was also significantly (*p* < 0.001) higher than that in bovine ETEC (< three plasmid replicons per isolate) (Fig. 5b). We further performed a genome-wide association study (GWAS) to determine the over- or under-representation of AMR genes or plasmid replicons in bovine and swine ETEC. Among the 95 types of AMR genes and 52 types of plasmid replicons detected in the 1,177 bovine and swine ETEC isolates, 25 types of AMR genes and 19 types of plasmid replicons were overrepresented in swine ETEC but underrepresented in bovine ETEC (Fig. 5c; Bonferroni corrected *p* value < 0.001). In contrast, only 2 types of AMR genes and 3 types of plasmid replicons were overrepresented in bovine ETEC but underrepresented in swine ETEC (Fig. 5c; Bonferroni corrected *p* value < 0.001). Our results indicated that the presence of more plasmid replicons might be associated with a higher likelihood of carrying multiple AMR genes in swine ETEC compared to bovine ETEC.

**Fig. 5:**
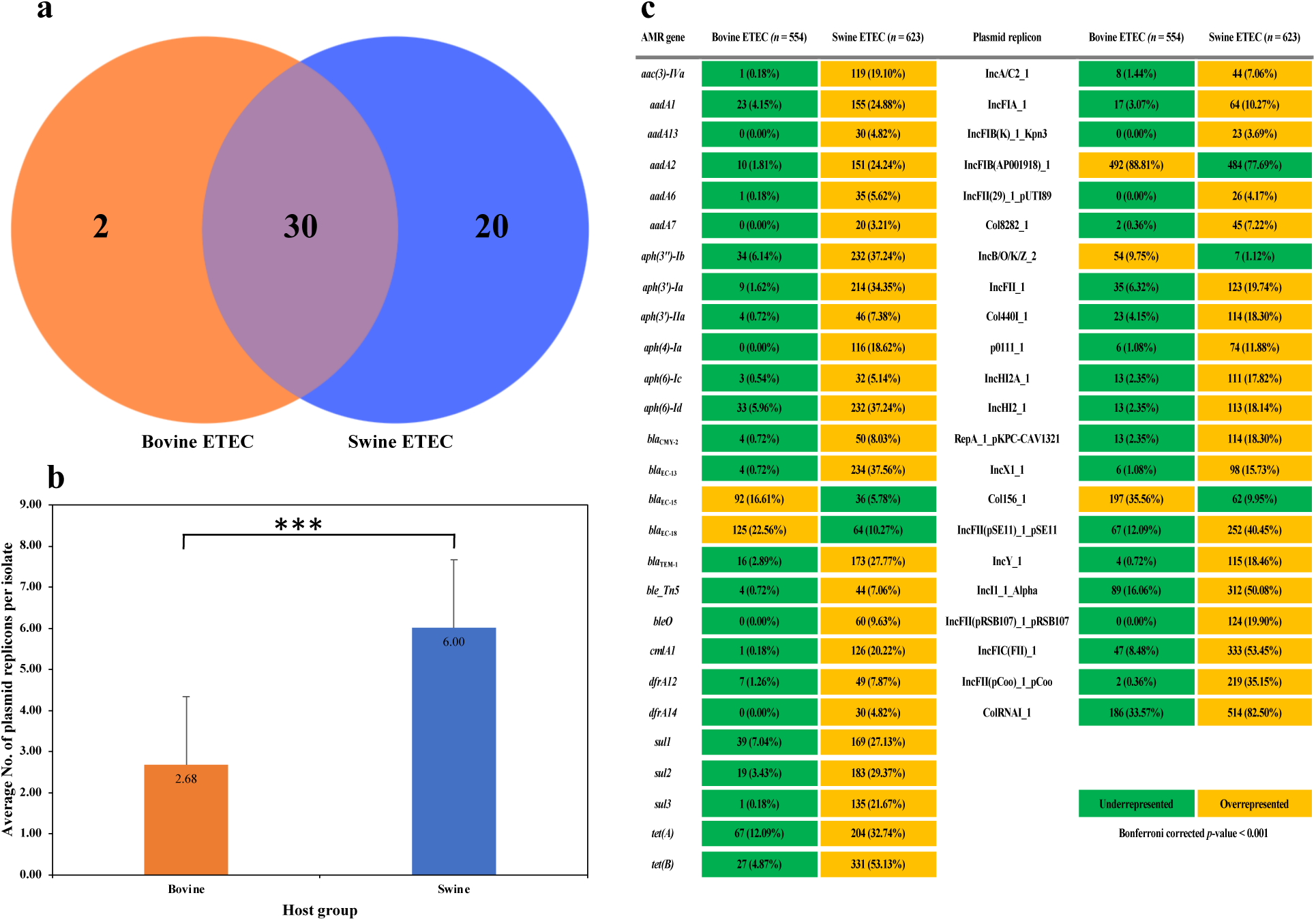
Positive correlation between abundance of plasmid replicons and prevalence of antimicrobial resistance (AMR) genes in bovine and swine ETEC. **a**, Number of plasmid replicons unique to or shared by bovine and swine ETEC. **b**, Average number of plasmid replicons per isolate detected in bovine and swine ETEC. **c**, Over- or under-represented plasmid replicons and AMR genes in bovine and swine ETEC. The *** in **b** indicates a significance level (*p* value) less than 0.001.

The overrepresented AMR genes in swine ETEC (Fig. 5c) encoded resistance to aminoglycoside [*aac(3)-Iva*, *aadA1*, *aadA2*, *aadA6*, *aadA7*, *aadA13*, *aph(3’’)-Ib*, *aph(3’)-Ia*, *aph(3’)-Iia*, *aph(4)-Ia*, *aph(6)-Ic*, and *aph(6)-Id*], beta-lactam (*bla*_CMY-2_, *bla*_EC-13_, *bla*_TEM-1_), bleomycin (*ble_Tn5*, *bleO*), chloramphenicol (*cmlA1*), trimethoprim (*dfrA12*, *dfrA14*), sulfonamide (*sul1*, *sul2*, *sul3*), and tetracycline [*tet*(A), *tet*(B)]. Most of these AMR genes are well documented plasmid-, transposon- or integron-encoded genes based on the Comprehensive Antibiotic Resistance Database (CARD). Further, the overrepresented plasmid replicons in swine ETEC (Fig. 5c) included those frequently associated with multi-AMR such as IncA/C_2_, IncHI2/IncHI2A, IncI1, IncX1^27, 28^. Although it was difficult to use the short-read sequence data to determine the precise location of AMR genes and associated mobile elements in this study, the above observation supported that there was a positive correlation between the presence of specific AMR genes and plasmid replicons.

### Higher prevalence of Shiga toxin genes in bovine ETEC than in swine ETEC

A key characteristic of ETEC is its ability to produce enterotoxins known as heat-labile (LT) and/or heat-stable (ST) toxins. Our virulence profiling (Supplementary Data 5) via Virulence Factor Database (VFDB)^29^ detected that 99.6% (552/554) of the bovine ETEC isolates carried the ST enterotoxin gene (*estIa*), only 0.4% (2/554) of the bovine ETEC isolates were positive for the LT enterotoxin genes (*eltA* and *eltB*) (Fig. 6a). On the other hand, 53.6% (334/623) and 40.0% (249/623) of the swine ETEC isolates were positive for the LT and ST enterotoxin genes, respectively (Fig. 6a). There were also 6.4% (40/623) of the swine ETEC isolates carrying both LT and ST enterotoxin genes, while the co-occurrence of the LT and ST enterotoxin genes was not detected in bovine ETEC isolates (Fig. 6a).

**Fig. 6:**
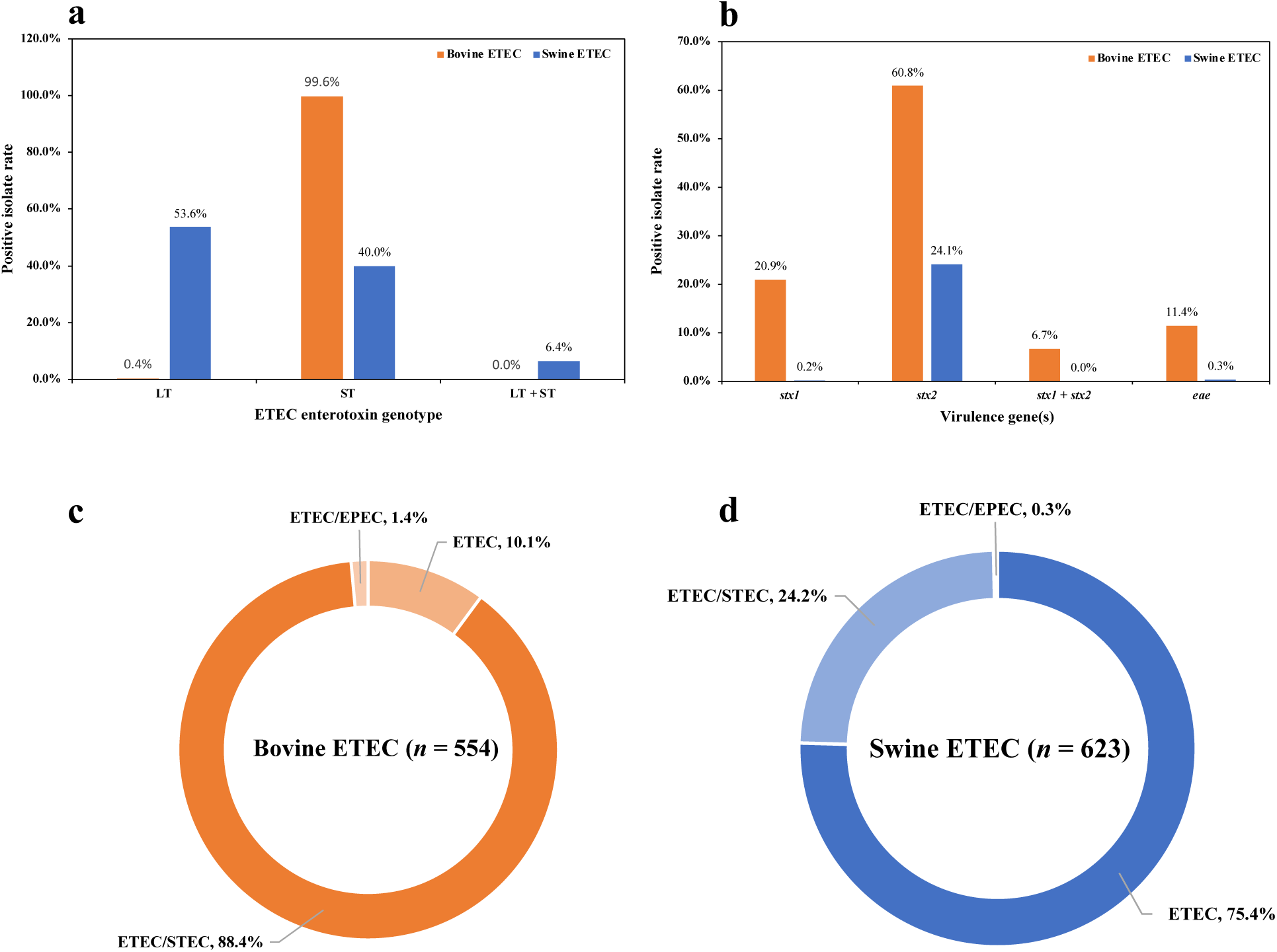
Co-occurrence of enterotoxin and Shiga toxin genes and the prevalence of hybrid pathovars in bovine and swine ETEC. **a**, Positive isolate rate for heat-labile (LT) and/or heat-stable (ST) toxin genes detected in bovine and swine ETEC. **b**, Positive isolate rate for Shiga toxin (*stx1*, *stx2*, *stx1* + *stx2*) and intimin (*eae*) genes detected in bovine and swine ETEC. **c**, Distribution of hybrid pathovar in bovine ETEC. **d**, Distribution of hybrid pathovar in swine ETEC.

In addition to enterotoxin genes, we also checked for other virulence factors such as Shiga toxin genes in the bovine and swine ETEC isolates to identify potential hybrid *E .coli* pathovars. Surprisingly, we found that the majority (490/554) of the bovine ETEC isolates harbored Shiga toxin genes (*stx1*: 20.9%, 116/554; *stx2*: 60.8%, 337/554; *stx1* + *stx2*: 6.7%, 37/554) (Fig. 6b), which are a defining feature of Shiga toxin-producing *E. coli* (STEC)^1^. Further, 11.4% (63/554) of the bovine ETEC isolates had an *eae* gene (Fig. 6b), an important virulence factor necessary for enteropathogenic *E. coli* (EPEC) or STEC to form attaching and effacing (A/E) lesions on epithelial cells^30, 31^. The presence of Shiga toxin or *eae* was much less common in swine ETEC than in bovine ETEC. Specifically, 24.1% (150/623) of the swine ETEC isolates were positive for *stx2* gene, while only 0.2% (1/623) were positive for *stx1*, and none of the swine isolates carried both *stx1* and *stx2* (Fig. 6b). Only 0.3% (2/623) of the swine ETEC isolates harbored an *eae* (Fig. 6b). We also investigated the subunits of *stx1* and *stx2* present in bovine and swine ETEC. In detail, all the *stx1*-positive bovine and swine ETEC isolates solely carried *stx1B* but lacked *stx1A*; however, almost all the *stx2*-positive ETEC isolates carried both *stx2A* and *stx2B*, except that *stx2A* was missing in one bovine ETEC isolate and *stx2B* was absent in two swine ETEC isolates. Based on the presence of specific virulence genes, we identified two types of potential hybrid *E. coli* pathovars (i.e., ETEC/STEC, ETEC/EPEC) in the bovine and swine isolates. The ETEC/STEC hybrid *E. coli* isolates accounted for 88.4% (490/554) of the bovine ETEC collection (Fig. 6c), but only 24.2% (151/623) of the swine ETEC collection (Fig. 6d).

## Discussion

Diarrhea caused by ETEC is one of the most common diseases in young calves and piglets, and investigation on the genotypes, AMR, and virulence profiles of livestock-associated ETEC is essential for guiding effective prevention and control strategies to mitigate the impact of ETEC on livestock. By analyzing the largest collection of bovine and swine ETEC isolates sampled over broad spatial and temporal scales, our study contributes significantly to the understanding of livestock-associated ETEC in the US. We found that distinct genotypes were associated with bovine and swine ETEC. Further, AMR patterns and virulence profiles in bovine and swine ETEC are quite different. Resistant ETEC isolates were more likely to be detected in pigs than in cattle, while hybrid pathovars (e.g., ETEC/STEC) with multiple virulence factors were more prevalent in cattle than in pigs. The observed genomic differences between bovine and swine ETEC in the US might be attributed to host adaptation and antibiotic usage practices.

Although more than 65% swine ETEC genomes and ∼89% bovine ETEC genomes deposited at EnteroBase are from the US, a systematic genomic analysis regarding the ETEC genotypes based on the sequencing data is missing in the literature. In this study, we performed WGS-based genotyping based on the ETEC genomes at EnteroBase. We observed that diverse genotypes were distributed among bovine and swine ETEC in the US. Specifically, we identified 17 *fimH* types, 21 O serogroups and 20 H serogroups in bovine ETEC, while 36 *fimH* types, 28 O serogroups and 32 H serogroups in swine ETEC. Moreover, a total of 36 and 85 STs were detected in bovine and swine ETEC, respectively (Supplementary Data 2). Clearly, swine ETEC is more diverse than bovine ETEC regarding the number of ETEC genotypes detected in each host group. Our observation is similar with previous studies from other countries that ETEC are highly diverse in terms of O and H serotypes^4, 6, 32^. The highly diverse ETEC genotypes detected in livestock is likely due to the plasmid-borne nature of the genes encoding ST and LT enterotoxins^33^. The plasmid-mediated transfer of enterotoxin genes may contribute to the diversity of ETEC strains as plasmids can move between bacterial cells, facilitating the exchange of genetic material. Although both bovine and swine ETEC show a great diversity in genotypes, distinct dominant genotypes are associated with each host species. Interestingly, the top three *fimH* types (*fimH*86, *fimH*444, *fimH*54), O serogroups (O168, O2, O136), H serogroups (H8, H12, H10) and STs (ST329, ST718, ST10) in bovine ETEC had little overlap with those present in swine ETEC (*fimH* types: *fimH*54, *fimH*24, *fimH*121; O serogroups: O149, O147, O8; H serogroups: H19, H4, H10; STs: ST90, ST10, ST100) (Supplementary Data 2). The spillover of ETEC genotypes from cattle to pigs or vice versa is seldomly detected in our study, indicating host adaptation may shape the genetic diversity and contributing to new variants of this pathovar in different food animals.

The use of antibiotics in various host environments can drive the evolution of resistance in pathogens. Different host populations may have varying levels of exposure to these agents^34^, which can influence the selection pressure for resistant strains. Historically, antibiotics have been used in both the swine and bovine industries for various purposes, including disease prevention, growth promotion, and treatment of infections^35^. However, the specific usage patterns and frequency can be influenced by factors such as industry practices, regulations, consumer demand, and veterinary recommendations^36^. For example, antibiotics are used more extensively in the swine industry compared to the bovine industry^34^. In addition, the types of antibiotic classes used in swine industry are also more diverse than those used in bovine industry; an exemplary antibiotic class is lincosamide (such as lincomycin and clindamycin), which is approved for use in pigs but not cattle^37^. The above facts may partially explain our results that swine ETEC had higher rate of isolates predicted to be resistant to the same antibiotic class and carried more diverse AMR genes than bovine ETEC. For instance, the percentage of isolates predicted to be resistant to common antibiotic classes used in livestock farming was 70.5% vs 10.8% for aminoglycoside, 62.3% vs 9.0% for sulfonamide, and 79.3% vs 19.7% for tetracycline in swine and bovine ETEC. Moreover, among the 95 types of AMR genes detected in the whole ETEC collection, 51 were unique to swine ETEC (e.g., AMR genes conferring resistance to lincosamide, fosfomycin, rifamycin) and only 4 were specific to bovine ETEC. The prevalence of AMR plasmids such as IncA/C_2_, IncHI2/IncHI2A, IncI1, and IncX1 in swine ETEC, combined with the intensive swine production system, can further facilitate the spread of AMR genes within the swine population or beyond. The difference in resistance rates and resistance gene diversity between swine and bovine ETEC emphasizes the need for tailored strategies to mitigate AMR in different food animals. More importantly, the prevalence of AMR genes conferring resistance to critically or highly important antimicrobials for human medicine (e.g., aminoglycoside, macrolide, fosfomycin, fluoroquinolone, chloramphenicol, lincosamide, sulfonamide, and tetracycline)^38^ in swine ETEC is concerning as these genes could be shared with foodborne pathogens through mechanisms like plasmid transfer. This could contribute to the growing problem of antimicrobial resistance, making infections harder to treat in both animals and humans.

Our virulence profiling revealed that a great number of bovine ETEC isolates were also identified as Shiga toxin-producing *E. coli* (STEC). STEC are a group of bacteria that can produce toxins known as Shiga toxins. This *E. coli* pathovar is often associated with foodborne outbreaks and can infect humans to lead to illnesses such as bloody diarrhea, and in severe cases hemolytic uremic syndrome (HUS), which can be life-threatening^39, 40^. Although STEC strains have been isolated from a variety of domestic and wild animals, certain host species are more significant in the maintenance and transmission of these bacteria. Cattle, in particular, have been recognized as the major reservoir host for STEC^41^. The fact that cattle serve as a major reservoir host for STEC sheds light on our observation that the detection rate of Shiga toxin genes was much higher in bovine ETEC compared to swine ETEC (Fig. 6b). The genes encoding Shiga toxins are often carried by lysogenic bacteriophages integrated in STEC chromosomes^42^. Theoretically, the Shiga toxin genes carried by bacteriophage can be transferred and integrated into the ETEC chromosome from the STEC chromosome via horizontal gene transfer (HGT), leading to the emergence of hybrid *E. coli* pathovar (i.e., ETEC/STEC). As the primary reservoir for STEC, cattle can provide a favorable host environment that facilitates HGT of virulence genes between ETEC and STEC strains. This aligns with our observation that the ETEC/STEC hybrid pathovars were more prevalent in cattle (88.4%; 490/554) than in pigs (24.2%; 151/623). The carriage of Shiga toxin-encoding prophage in an ETEC isolate may provide a selective advantage in cattle^43, 44^. Nevertheless, the emergence of hybrid pathovars of *E. coli* can have significant implications for public health, agriculture, and food safety as the hybrids potentially have increased virulence or enhanced disease-causing abilities, broader host range, and altered transmission patterns^45^.

A limitation of this study is our AMR profiling relies solely on genotypic prediction and lacks phenotypic confirmation. Genotypic prediction of AMR is based on known resistance genes in the bacterial genomes. Not all resistance mechanisms are well understood, and some genes may not yet be identified or linked to resistance. Reliance on known gene markers could miss emerging or novel resistance mechanisms. Further, genotypic prediction alone might not accurately reflect the potential for resistance. For example, some resistance genes such as the *blac*_EC_ family genes detected in this study might have no effect on phenotypic resistance^22^. To overcome this limitation, studies should ideally combine genotypic prediction with phenotypic testing to confirm actual resistance. Another limitation of our study is although we observe a strong correlation between plasmid replicon content and AMR gene prevalence based on GWAS analysis, it is difficult to accurately locate AMR genes on plasmids using short-read sequence data in the study. Long-read sequencing technology should be employed to obtain the complete plasmid sequences and identify the specific AMR genes associated with them.

In conclusion, our study unraveled the dissimilarity in genotype, AMR and virulence between bovine and swine ETEC in the US. Understanding these differences can aid in the development of targeted preventive and therapeutic strategies for controlling ETEC-related gastrointestinal illnesses in both livestock and humans. In addition, understanding the role of host ecology in shaping ETEC diversity and characteristics can also provide insights into the broader context of bacterial evolution and host-pathogen interactions. Finally, as bacteria like ETEC can undergo rapid evolution in different host species and agricultural practices such as antibiotic usage are subject to change over time in livestock farming, ongoing surveillance and research are essential to monitor the genetic dynamics of ETEC in both cattle and pigs.

## Methods

### Dataset collection

ETEC genomes from bovine and swine hosts were retrieved from EnteroBase on March 10, 2023 (Supplementary Data 1) using the following search terms: species-*Escherichia coli*; source niche-livestock; source type-swine or bovine; predicted pathovar-ETEC; country-United States. A total of 1,264 US bovine and swine ETEC genomes were deposited at EnteroBase as of the retrieval time. We filtered the ETEC genomes that were not accessible due to delayed release time settings, had low read quality (See **Method-Quaity assessment for raw reads**), or lacked exact collection location (i.e., US states). The refined collection consisted of 1,177 ETEC genomes. The ETEC isolates were collected over broad spatial and temporal scales (Fig. 1), including 554 bovine ETEC isolates collected from 28 US states during 1976-2023 (Fig. 1a), and 623 swine ETEC isolates collected from 35 US states during 1970-2023. Detailed metadata information (isolate name, US state, collection year, biosample accession number, SRA accession number, BioProject number, etc.) of each isolate in the collection was provided in Supplementary Data 1.

### DNA extraction and whole-genome sequencing

For DNA extraction of the ETEC isolates, each isolate was streaked onto MacConkey agar plates and incubated for 18 h at 37 °C. A single colony was then picked, transferred to Luria-Bertani broth, and cultured overnight at 37 °C with continuous agitation (250 rpm). Genomic DNA was extracted using the Qiagen Dneasy^®^ Blood & Tissue kit (Qiagen, Valencia, CA, US) following the manufacturer’s instructions. DNA purity (1.8 ≤ A260/A280 ≤ 2.0) was confirmed using NanoDrop™ One (Thermo Scientific™, DE, US) and DNA concentration was quantified using Qubit^®^ 3.0 fluorometer (Thermo Fisher Scientific Inc., MA, US). Extracted genomic DNA was stored at -20 °C before WGS. For WGS, DNA library was prepared using the Nextera XT DNA Library Prep Kit (Illumina Inc., San Diego, CA, US), normalized using quantitation-based procedure, and pooled together at equal volume. The pooled library (600 μL at 20 picomolar) was denatured and sequenced on an Illumina MiSeq sequencer (Illumina Inc., San Diego, CA, US).

### Quality assessment for raw reads

The quality of the raw reads obtained in this study and downloaded from EnteroBase was assessed using the MicroRunQC workflow in GalaxyTrakr v2^46^. Sequence data passing quality control thresholds (i.e., average coverage ≥ 40, average quality score ≥ 30, total assembly length between 4.5 and 5.9 Mb) were used for subsequent genomic analyses.

### Whole-genome sequencing-based genotyping

Raw reads of the bovine and swine ETEC isolates were *de novo* assembled using Shovill (Galaxy v1.0.4) to obtain the draft genome assembly^47^. With the draft genomes of the ETEC isolates as input, ClermonTyper^15^ was used to determine Clermont phylogroups, FimTyper^16^ for *fimH* types, and ECTyper^19^ in combination with EtoKi EBEis (EnteroBase *Escherichia in silico* serotyping module from EnteroBase Tool Kit)^20^ for serotype prediction (O and H serogroups). Sequence types (ST) of the ETEC isolates were identified using classic seven-gene (*adk*, *fumC*, *gyrB*, *icd*, *mdh*, *purA*, and *recA*) multilocus sequence typing (MLST) scheme^18^ at EnteroBase.

### AMR, plasmid replicon, and virulence profiling

ABRicate (Galaxy v1.0.1)^48^ was used to identify the AMR genes, plasmid replicons, and virulence factors by aligning each draft genome assembly against the NCBI AMRFinder database^21^, PlasmidFinder database^26^, and Virulence Factor Database (VFDB)^29^, respectively. The default settings of ABRicate (i.e., minimum nucleotide identity and coverage thresholds of 80% and 80%) were used for all searches.

### Detection of point mutation

The draft genome assembly obtained previously were aligned against the PointFinder database^24^ to identify chromosomal mutations mediating antimicrobial resistance. We identified chromosomal point mutations in *ampC* promoter (beta-lactam), *pmrB* (colistin), *gyrA* or *parC* (fluoroquinolone), and *folP* (sulfonamide). The point mutations identified using PointFinder in combination with the AMR genes detected using NCBI AMRFinder (Supplementary Data 3) were integrated for AMR profiling.

### Genome-wide association study of the over- and under-representation of AMR genes and plasmid replicons

The presence and absence of AMR genes or plasmid replicons identified by Abricate were used for Scoary analyses. For each strain in our dataset, the presence of a given trait in this strain (i.e., an AMR gene or plasmid replicon) was indicated by a value of 1 and the absence of a trait by a value of 0. The resulting presence-absence matrices, one for AMR genes and one for plasmid replicons, were provided to Scoary v1.6.16^49^ as “gene” input tables. A second file assigning strains to either the swine or bovine category was used as the “trait” table for each input. Genes in each matrix with identical distributions were consolidated with the –collapse flag, and the output was limited to associations with a naïve *p*-value ≤ 0.05. Associations with a Bonferroni-corrected *p*-value ≤ 0.001 were chosen for further analysis.

### Statistical analysis

The average number of AMR genes per isolate and the average number of plasmid replicons per isolate carried by bovine ETEC and swine ETEC, together with their standard deviations were calculated. Differences in average number of AMR genes per isolate as well as difference in average number of plasmid replicons per isolate between bovine ETEC and swine ETEC were assessed using student’s *t*-test, and were considered to be significant when *p* < 0.001 (SAS 9.4, SAS Institute Inc., Cary, NC, US).

## Data availability

Sequence data of the ETEC isolates from our lab are deposited in the NCBI Sequence Read Archive (SRA) (https://www.ncbi.nlm.nih.gov/sra) under BioProject <u>PRJNA357722</u>. Publicly available sequence data are downloaded from EnteroBase (https://enterobase.warwick.ac.uk/), NCBI SRA (https://www.ncbi.nlm.nih.gov/sra) and the European Nucleotide Archive (https://www.ebi.ac.uk/ena). Accession numbers of the genomes used in this study are listed in Supplementary Data 1.

## Acknowledgements

This work is supported by the US Food and Drug Administration (Grant No. 1U19FD007114-01), US Department of Agriculture (Grant No. PEN4522), and Penn State College of Agricultural Sciences.

## Contributions

Y.F. designed the study, sequenced part of the ETEC collection and retrieved the genomes of the ETEC collection from EnteroBase, performed the majority of bioinformatics analyses of the data, interpreted the data, and wrote the draft manuscript; E.M.N analyzed the enrichment of genes in bovine and swine ETEC using Scoary. N.M.M. and E.G.D. contributed to interpretation of the data and manuscript revision.

## Corresponding author

Correspondence to Yezhi Fu and Edward G. Dudley.

## Ethics declarations

The authors declare no competing interests.

